# The first one hundred nanometers inside the pre-synaptic terminal where calcium diffusion triggers vesicular release

**DOI:** 10.1101/326835

**Authors:** C. Guerrier, D. Holcman

## Abstract

Calcium diffusion in the thin one hundred nanometers layer located between the plasma membrane and docked vesicles in the pre-synaptic terminal of neuronal cells mediates vesicular fusion and synaptic transmission. Accounting for the narrow-cusp geometry located underneath the vesicle is a key ingredient that defines the probability and the time scale of calcium diffusion to bind calcium sensors for the initiation of vesicular release. We study here the time scale, the calcium binding dynamics and the consequences for asynchronous versus synchronous release. To conclude, threedimensional modeling approaches and the associated coarse-grained simulations can now account efficiently for the precise co-organization of vesicles and Voltage-Gated-Calcium-Channel (VGCC). This co-organization is a key determinant of short-term plasticity and it shapes asynchronous release. Moreover, changing the location of VGCC from few nanometers underneath the vesicle modifies significantly the release probability. Finally, by modifying the calcium buffer concentration, a single synapse can switch from facilitation to depression.

## Introduction

The first one hundred nanometer domain, between the plasma membrane and the vesicles in the pre-synaptic terminal remains difficult to study, yet it seems that a displacement as small as a ten of nanometers in the molecular organization can affect vesicular release. There are many examples, where a ten nanometer precision has to be achieved in order to guarantee normal physiology function. This is the case for the apposition of pre- and post-synaptic terminal of neuronal synapses: this apposition is obtained by a set of redundant adhesion molecules, such as laminins that self-organize to maintain the synapse structure and stability in the central nervous system. A lack of the laminin *β*_2_ subunit leads to a disruption of the hippocampal synapse structure, to a misalignment of the pre- and post-synaptic partners and to an increased post-synaptic density (PSD) size [1]. In addition, mutations in PSD proteins are associated with neurological and psychiatric diseases [2]. Another example is autism spectrum disorders which have been associated with the mutations in genes encoding Shank2 and Shank3, PSD-93, and a mis-regulation of adhesion molecules neuroligin 3, neuroligin 4, and neurexin 1 [3, 2], affecting a precise geometrical apposition.

In the pre-synaptic terminal, some vesicles are concentrated in a region called the Active Zone (AZ), which is well aligned with the PSD of the post-synaptic terminal. This apposition creates geometrical columns: a single column alignment was originally hypothesized and numerical simulations showed that it maximizes the synaptic current [4] while minimizing its fluctuations [5]. Multiple nanocolumns were predicted in [6] to sustain synaptic response and a more reliable transmission compared to several synapses containing a single column. Finally, these columns have recently been confirmed experimentally and observed at superresolution [7, 8]. This nanocolumn example shows that synaptic transmission uses a tens of nanometer precision for its organization and for example a misalignment of synaptic terminals is at the basis of several pathological disorders [9].

Another example is the PSD, that cannot permanently retain glutamatergic receptors that are moving by random motion [10]. After a long enough time, these receptors spread out, modifying the synaptic current [6, 11]. In the absence of direct experimental approaches, studying the functional consequences of the nanometer precision in the domain between the membrane and the vesicles at the pre-synaptic terminal has recently benefited from three-dimensional modeling and numerical simulations. We focus this manuscript on this paradigm shift of analyzing the diffusion of calcium ions and in particular about the nanometric relation between the organization of calcium channels and vesicles and how it shapes the release probability, synaptic transmission, asynchronous release and short-term plasticity.

## 1 Dynamics and constant re-organization in the pre-synaptic terminal

Despite the fast advances of super-resolution microscopy, that allowed to reconstruct structural *in vivo* cell properties, or to follow calcium and voltage using genetically encoded indicators [12, 13, 14, 15], it remains difficult to study the detailed molecular dynamics in nanometer domains at a time scale less than 100 ms. Indeed, to understand how molecules interact in nanometer domains, the notion of concentration has to be abandoned because it does not make much sense due to the large fluctuations in the small number of molecules. However, molecular interactions can still be transformed into a cellular activation at the micrometer level, but the exact biophysical mechanisms remain in most cases unclear or controversial. Modeling and numerical simulations based on biophysical principles have emerged as orthogonal tools compared to experiments to describe molecular dynamics at this spatio-temporal scales [16].

At this intermediate level between the molecular and the cellular scale, physical modeling of diffusion is based on Brownian motion, which requires to specify an inherent time scale of simulations. Indeed the motion of molecules follows a random walk approximation expressed by the Euler’s scheme for a trajectory ***X*** (*t*) at time *t*:

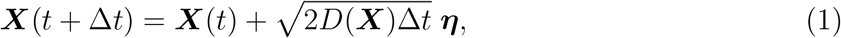

where ***η*** is a Gaussian variable, *D*(***X***) is the spatial dependent diffusion coefficient and Δ*t* is the time scale to be chosen. It is usually a difficult choice. It should not be too small to avoid wasting simulation times and should not be too large compared to the small spatial scales involved in the microdomain, such as molecular binding sites.

In particular, taking into account in numerical simulations the region between vesicles and the plasma membrane has been particularly difficult to model because of its cusp-like geometry. It requires a specific mathematical treatment to estimate the mean time for a calcium ion after entering through a Voltage-Gated-Calcium-Channel (VGCC) to find a key calcium binding sensor, involved in triggering vesicular release [17] (Fig. 1). Such sites are Ca^2+^- binding proteins, located on synaptotagmins, that are involved in triggering directly or not vesicular fusion [18]. They are located precisely in this nanometric region below vesicles. Calcium diffusion in the pre-synaptic terminal has traditionally been modeled as two-or three-dimensional diffusion [19,20, 21], but ignoring the three-dimensional complications of the vesicle shape. However, for auditory hair cells, Monte-Carlo simulations revealed [22] that the spherical shape of the ribbon where vesicles are tethered, can generate a local Ca^2+^ microdomain that enhances vesicular fusion by trapping calcium ions [22]. This spherical ribbon that aggregates vesicles is likely to create an intermediate microdomain for calcium dynamics between the pre-synaptic bulk and the boundary layer near the membrane, which should be further investigated. For other types of synapses, while estimating the time scale of calcium binding, and computing the vesicular release probability, one cannot ignore the specific three-dimensional organization of the first one hundred nanometer, the region underneath vesicles and the position of calcium sensors. The release probability not only depends on the binding of calcium ions to sensor proteins, located underneath the vesicle [23], but also on the AZ organization: a sparse vesicular distribution vs vesicular crowding, and channels clustered vs uniformly distributed (Fig. 1B)[24, 25]. The major components in these dependencies being the distance between VGCC and vesicles coupled to the particular cusp-like geometry [23].

**Figure 1:**
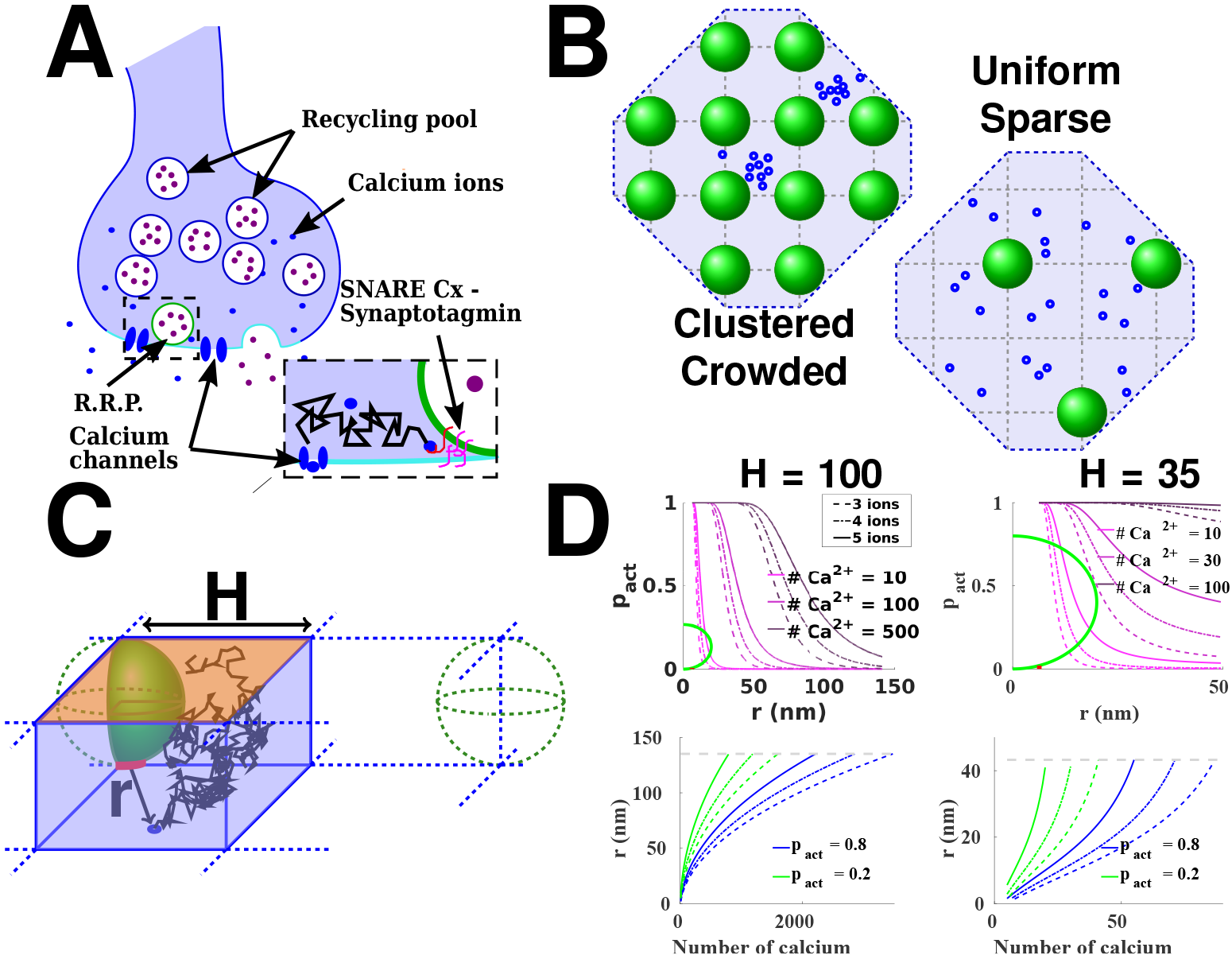
Estimating the release probability. **A**: Functional organization of the presynaptic terminal. An incoming action potential leads to the opening of voltage gated calcium channels (blue) located at the AZ (light blue). The consecutive entry of calcium ions (orange) triggers the fusion of docked vesicles (green) with the synaptic membrane, and the liberation of neurotransmitters (purple) in the synaptic cleft. The binding of neurotransmitters to specific receptors located in the post-synaptic terminal triggers the conversion of the chemical signal into an electrical signal in the post-synaptic neuron. **B**: Model of the AZ organization. Vesicles (green) are regularly (left) or sparsely (right) distributed on a square lattice. Calcium channels (blue) can be clustered (left) or uniformly distributed (right) in the AZ. **C**: Elementary 3-dimensional domain to compute the splitting probability for an ion starting in the bottom of the domain (blue circle representing a channel), to reach the target (red) before leaving the domain through the orange boundary. The other boundaries are reflecting. The vesicles are distributed on a square lattice of side 2*H*. *D*: Top: Probability to find three, four or five calcium ions (full, dashed and dotted lines respectively) underneath a vesicle, in the case of sparse vesicular distribution: *H* =100 nm (left) and in the case of crowding of vesicles at the AZ: *H* =35 nm (right). The relation depends on the initial number of calcium ions. The diameter of the pre-synaptic vesicles is fixed at R=40 nm (green), the diffusion coefficient for free calcium ions being *D*_Ca_ = 200*μm*^2^*s*^−1^. Bottom: Maximal channels distance *r* to activate the vesicle with a probability *p*_*act*_ ≥ 0.8 (blue) and 0.2 (green), when there are *N* initial ions, for *H* = 100 nm (left) and *H* = 35 nm (right). We fix the threshold to 3, 4 or 5 calcium ions. The gray dashed line represents the maximal distance to the vesicle in the elementary domain: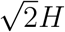.

## 2 Calcium binding sensors and vesicular release kinetics

The synaptotagmin family of molecules are Ca^2+^ sensors for vesicle fusion: following Ca^2+^ binding, activation of the SNARE-complex mediates membrane fusion. Although there are two specific sensors on synaptotagmin, the number of ions necessary for fusion was estimated around 4 or 5 [26, 27]. This apparent discrepancy comes from the indirect method of analysis, that may also account for calcium bound to calcium sensor buffers.

### 2.1 Modeling calcium binding and limitation of using a forward rate constant

Modeling the causality between calcium dynamics and vesicular fusion relies on patching steps, resulting in simulations that do not necessarily account for the three-dimensional organization of the AZ. In [28], by using the software Mcell, the forward rate constant of calcium ions is the reciprocal of the flux to the sensor targets is assumed rather than derived from physical considerations. This rate has to be pre-calculated especially when calcium sensors are located underneath vesicles (see next section). In addition, due to the vesicle crowding, the calcium flow cannot be represented by a Gaussian function, which is the classical probability density function of independent particles initially concentrated at one spot in a free space. This approximation further neglects the boundary effect and the depth of a synapse. In that context, providing numbers such as a distance of 30 nm for a possible exclusion area between vesicles and VGCCs, based on a two-dimensional approximation that ignores the effect of the vesicle size with a radius of *R* = 20 nm is not necessarily accurate. But the question is by how much? Clearly this geometrical limitation calls for a three-dimensional approach accounting for vesicular structure and organization.

A different modeling approach is described in [25], based on diffusion [29, 30] of a twodimensional coarse-grained lattice, using a two-dimensional Gaussian approximation for the initial calcium entrance. The vesicular release in these approaches is computed by using a Markov model, which is based on the concentration of calcium in a two-dimensional domain around the location of the channel. However, this concentration does fluctuate a lot, depending on the size of the sampling volume. This modeling approach neglects the small number of ions that penetrate underneath the vesicle, which is replaced by assuming a value for the forward rate. Indeed, to trigger vesicular release from the binding of calcium ions to sensors, a value should be given to the forward rate *k*_*on*_. Since the early work of von Smoluchowski in 1916 [31], this rate has been computed as the flux from a fixed concentration to a narrow window located on an infinite plan. But these assumptions are not satisfied for the small steady-state calcium concentration, because the rate should be computed for the first calcium arriving to the binding site, sampled from the transient entrance through the VGCC. In summary, using a forward rate constant presuppose a geometrical organization. An alternative is to replace such a rate by Brownian simulations, but this approach is in general heavy computationally. Recent hybrid simulations have been developed, where classical diffusion is used far away from a sensor, while near the boundary of a channel a Brownian representation is used (see [32] for a description of such framework).

### 2.2 How to chose an effective Diffusion coefficient

What should be the buffer distribution and concentration in the pre-synaptic terminal? Various buffers and concentrations were previously considered: for the calyx of Held, a concentration of 400 *μM* immobile endogenous buffers plus a Parvalbumin-like Ca^2+^ diffusing buffer with concentration of 50 *μM* were used in [28], while ATP, a mobile Ca^2+^ buffer, was present at a total concentration of 2 mM in all simulations (with a diffusion coefficient of 220 *μm*^2^/*s*). In [24] the concentration of fixed buffers is 480 *μM*, and of mobile buffers is 100 *μM* with *D* = 20 *μm*^2^/*s*. The buffer capacity *κ*_*S*_(ratio of bound vs free) is often chosen equal to 40. However, one of the main free parameter remaining is the value of the forward rate, which is valid for a single compartment model like the pre-synaptic terminal, which disregard its heterogeneity, but not for buffers located underneath vesicles.

### 2.3 Calcium-Buffer interactions in the pre-synaptic bulk

When the pre-synaptic terminal is modeled as a bulk only, it does not matter that buffers are moving or not, because in one compartment the differential equations disregard the geometry. So, what matters is the number of free buffers available. If buffers are modeled with stochastic simulations, then space matters, especially during multiple entry of calcium channels, due to fluctuations of calcium buffers in the region very close to the calcium sensor sites underneath the vesicle. But to be efficient, this geometry should be implemented, which is often difficult.

In summary, buffers could be homogeneously distributed in the bulk, but between vesicles, the concentration is much less homogenous. More drastically, the number of buffers between the membrane and the first layer of vesicles can be of the order of a few: indeed, for vesicles positioned on a square lattice with radius 60*nm* with a height of 40*nm*, the volume of the parallelepiped lattice *P*_*para*_ is *V* = 0.06 × 0.06 × 0.04 = 24 × 6 × 10^−6^, minus the volume of a vesicle which is 32 × 10^−6^*μm*^3^, that is Vol_total_ = 1.12 × 10^−4^*μm*^3^. Inside such a region, for a buffer concentration of 40 *μM*, this represents around 26 buffers. For 400 *μM* (at the calyx of Held), this represents around 260 buffers. These numbers should be compared to the number of free calcium entry (from 80 to 500). It is conceivable that the different vesicular proteins located near the vesicular calcium sensor (others than the synaptotagmins) play a more important role for buffering calcium than the diffusing calcium buffers located in the bulk which can occasionally enter into the region *P*_*para*_, because they are precisely located at the right place and thus could create an efficient local calcium reservoir.

Finally, calcium mitochondria uptake can affect synaptic release through the MCU channels
[33]. Mitochondrias participate in the calcium regulation that controls synaptic release and a MCU disruption could increase asynchronous release, decreasing the efficacy of synchronous neurotransmitter release and could also alter short-term presynaptic plasticity. This suggests that the distribution of mitochondria within the AZ could be as determinant as calcium buffers, a question that should be further investigated.

Another reason to reconsider the role of calcium buffer in the first one hundred nanometer layer is the presence of an electric field that could push ions inside the bulk, as revealed recently for the synaptic terminal [15, 34]. Too many buffers should disrupt calcium signalling and direct vesicular release. A low buffer capacity will increase asynchronous release. Thus, the concentration of buffers that favor a synchronous release should have an optimal value: not too low and not too large.

In [23, 35], the number of calcium entering through the VGCC vary from 80 to 500, which is also the case in [24], where they open around 12 Ca^2+^ channels with a single channel current of 0.15 pA and a duration of 105 *μs*, leading to ≈ 500 – 600 calcium ions. Some of the 260 buffers with at least two binding sites, could bind calcium ions on their direct way to the calcium sensor underneath the vesicle. With ten times less buffers, much more free calcium would be available and then what would matter is the distance of the channel to the vesicle.

### 2.4 Phenomenological laws between probability and the overall calcium concentration

The relation between the molecular organization of VGCC, their numbers, calcium buffer dynamics, the release probability *P*_*release*_ and the calcium flux of entering concentration of calcium *[Ca]*_*flux*_ mediated by an action potential remains an interesting problem. Over the
years the following empirical relation has been proposed [26]:

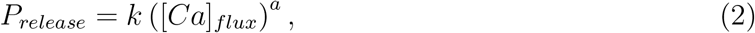

where *k* is a constant. The relation between the exponent *a* and the number of active sensor binding sites is not direct, due to the effect of buffers, the clustering of VGCCs, but also the local vesicular geometry. Indeed, the contribution of geometry appears while computing the probability for an ion to go underneath the vesicle vs going directly to the bulk [17, 28].

### 2.5 Partial conclusion: modeling vesicular release

To conclude, there is not yet a derived formula from physical principle to connect the flux or transient calcium concentration and the release probability, however, stochastic approaches are used to estimate the arrival of ions to the calcium sensors [25, 24, 23, 35]. In addition, the three dimensional vesicular organization should be accounted either directly, by implementing vesicles as obstacles or by computing the Brownian flux to small targets located underneath.

Following calcium diffusion, once calcium ions are bound to buffers, they can possibly unbind, but often the exact value of the backward rate constant is unknown. In recent mathematical models [28, 23, 35], vesicles are released when all 5 binding sites at a single sensor are occupied. If less than 5 calcium ions are bound, the vesicle is waiting for the final ions to arrive. There can be several copies of molecular sensors, but a single one might be sufficient to trigger release. It might also be conceivable that multiple binding sensors cooperate in the release process, and this possibility could explain the large modulation of the vesicular release probability [26, 27].

We already emphasised that the calcium ions bound to calcium sensors located underneath the vesicles can contribute critically to the residual calcium ions pool, especially when the backward rate is very small. To conclude, there are two types of calcium ions contributing to shaping the vesicular release probability. The ions already bound to the specific calcium sensors located underneath the vesicles, and the ions freely moving in the pre-synaptic terminal, that can reach the calcium sensors or induce calcium release from organelles, hence filling a binding site, and ultimately triggering vesicular fusion. This mechanism represents a possible scenario for the calcium contribution to the asynchronous vesicular release (see below).

## 3 Distribution of calcium ions entering through a VGCC

The Hodgkin-Huxley model [36] can be used [35] to generate a calcium influx current inside the pre-synaptic terminal. Following the opening of VGCC, this current corresponds to an entry per channel during a mean time of ≈1 ms for approximately 80 calcium ions, compatible with a previous estimation of 200 reviewed in [21] or with the 45 ions per channels described in [28]: with a total of 12 channels, this would represent 540 ions entering during ≈ 0.1 ms. After entry, the calcium flux can be split into a ionic component that reaches the small region below the vesicle, and another one reaching the pre-synaptic bulk. The probability for a calcium ion to reach the binding region, defined as a small ribbon joining the vesicle and the plasma membrane, before the bulk, has been computed in models where vesicles are organized in a square lattice (Fig. 1B) with length 2*H*, where *H* is of the order of the diameter of a vesicle, from 40 to 100 nm (Fig. 1C). For a dense set of vesicles distributed on a square lattice, the splitting probability for a calcium ion (modeled as Brownian) to reach the ribbon before the bulk is [23]:

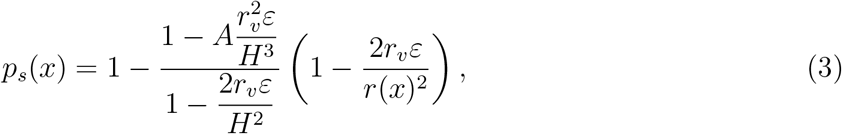

where *r*_υ_ is the size of a vesicle, *H* is half the distance between two vesicles, *A* = 9.8, *r*(*x*) is the distance between the point source and the closest vesicle, and ε is the height of the small cylindrical ribbon (fig. 1C), where calcium sensors are located. This probability accounts for the particular geometry of the target and depends on the relative distance between the targets and the source points [23].

### 3.1 Calcium time scales to the ribbon region

How long does it take for a calcium ion in the synaptic bulk or at the mouth of a VGCC (located far away from a vesicle) to enter into the cylindrical ribbon (red region in Fig. 1A-C) underneath a vesicle? This mean time computed analytically in [23, 16] is given by:

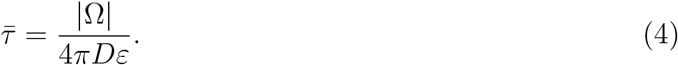

For a volume of a pre-synaptic microdomain |Ω| = 1 *μm*^3^, a diffusion coefficient *D* = 20 *μm^2^*/*s* and a size of the ribbon in the range ε = 0.001 – 0.01 *μm,* the mean time is 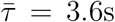. We note that the diffusion coefficient of calcium ions in a free environment is usually *D* ≈ 200*μm*^2^/*s*, but the motion of ions in pre-synaptic terminals or in dendrites is restricted by obstacles such as microtubules, actin and organelles. In [37], the effects of crowding on the diffusion coefficient has been estimated using modeling, simulations and a cytoplasmic fluid in a patch pipette, leading to a modified effective diffusion coefficient *D* ≈ 20*μm*^2^/*s*.

However the size of a calcium ion of the order of 1*nm* should not affect the classical law of diffusion. So the size of a calcium ion is often neglected in most modeling and stochastic simulations. Certainly, the most interesting part of vesicular crowding micro-environments is the local molecular organization underneath the vesicle, formed by all vesicular molecules, such as SNARE, syntaxin or synaptotagmin, that could result in a ten to twenty-nanometer environment filled with polymer filaments. This intermediate spatial scale could have several effects such as 1) sequestrating calcium ions and/or creating channels to any sensor sites, 2) preventing calcium channels to get too close to the calcium sensors 3) positioning synaptotagmin close to VGCC.

To conclude, as crowding is the main obstacle to diffusion, when the diffusion time scale
involves long distances the effective diffusion coefficient should be used, while for short-distances containing little obstacles, computation should be performed with the cytoplasmic diffusion coefficient. No evidences have shown that changes of the cytoplasmic volumes, occurring at a time scale of milliseconds can modify the nature of calcium diffusion.

### 3.2 Direct and indirect vesicular release activation

Using the values mentioned in the previous subsection, the mean time for calcium to transit from the bulk underneath a vesicle is 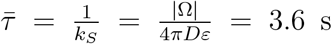 (for ɛ =1 nm) to 360ms (for 10 nm). This mean time is much longer than the initial calcium transient from the channels(< 5 *μs*), as we shall see now. The reciprocal of this time is the Poissonian rate *k*_*S*_ representing the rate of arrival of a free calcium ion to a binding sensor. However, this time is very different from the time for an ion entering through a VGCC close to the vesicle, to reach the region underneath the vesicle directly, i.e. while staying in a boundary layer around it. Indeed due to the confinement by the vesicle and in the absence of large obstacles at a distance of 20 nm, the time for an ion to hit a target sensor is 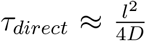 Using the free diffusion coefficient for calcium *D* ≈ 200 *μm*^2^/*s*, this leads to a mean time of 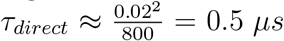. It is however possible to observe faster activation events: indeed vesicular release can be activated by the fastest ions arriving to the sensor site, in that case, if there are *N* calcium ions entering through a channel, then the fastest arrival of two ions falls into the extreme statistics analysis and the mean time is 2 * *τ*_*direct*_/*log*(*N*) [38]. For *N* = 100, we get around 0.0005 *μs*. This scenario provides a physical mechanism for the fastest transmission events reported in [39].

## 4 Computing the release probability when VGCC are located underneath a vesicle

The probability that a finite number *T* of calcium ions (*T* =3,4 and 5 ions) are bound at specific binding sites located between a vesicle and the synaptic membrane (subsection 3.1), when *N* calcium ions have entered through a cluster or a single VGCC located at a distance *r* from the center of the closest vesicle is defined by:

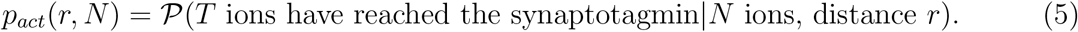

When the calcium unbinding events are too slow to be taken into account, the probability of vesicular release *p*_*act*_(*r*, *N*) is thus the one to find at least *T* ions inside the cylindrical ribbon. The probability to find exactly *k* ions out of *N* follows the Binomial distribution ***B***(*N*,*p*_*s*_(*r*)), and the steady-state probability is:

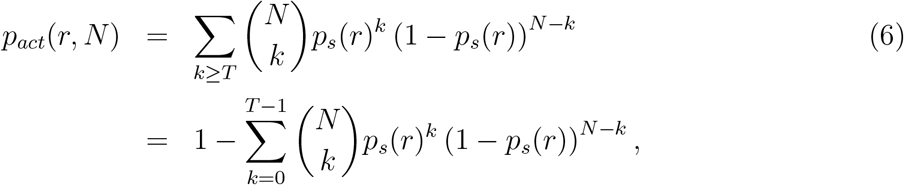

where *p*_*s*_(*r*) was computed in Eq. 3. The maximal VGCC distance *r*_*max,pact*_(*N*) to activate a vesicle with a probability *p*_*act*_ ≥ 0.8 is shown in Fig. 1D. The probability *p*_*act*_ critically depends on the channels-vesicles distance, which can vary from a few to hundreds of nanometers. This fast decrease of the probability with the distance explains the large variability in the release probability as VGCC position can vary over time [40].

The organization of vesicles in the AZ also influences the release probability: when vesicles are sparsely distributed (Fig. 1B-D, *H* = 100 nm) and 100 ions entered through VGCC, then a 80% release probability *p*_*act*_ = 0.8 is reached when the distance between the vesicles and the channels is smaller than 24 nm, compared to the 20 nm radius of the vesicle. This result shows that the co-localization of VGCC with a vesicle is a key feature determining a high synchronous release probability. However, for a high vesicular crowding (Fig. 1B-D, described by choosing the distance *H* = 35 nm) and when 100 ions are released instantaneously at a VGCC, then the probability *p*_*act*_ is higher than 0.9, regardless of the initial position of the channel, suggesting that vesicles are certainly released and leading to a synchronous release.

To conclude, a high crowding of vesicles should be associated with a high-release probability sustaining a synchronous release, while a sparse vesicle density might be associated with asynchronous release. Channels can be organized in clusters or uniformly distributed and this is also a major determinant governing release probability (Fig. 1B-D). Indeed, the effect of channels clustering can be modeled by simply increasing the number of entering calcium ions. When vesicles are sparsely distributed, the 24 nm distance required to obtain a release probability *p*_*act*_ = 0.8 when 100 ions are entering through one channel, is increased to 61nm for 500 ions. This effect results from the local geometry of the ribbon underneath the vesicle. When the number of ions is low, this maximal distance to guarantee *p*_*act*_ = 0.8 does not vary much when the activation threshold *T* increases from 3 to 5; however, for 500 ions, this distance changes significantly over 15 nm.

The maximal distance *r_max,Pact_ (N*) between channels and vesicles to obtain a given release probability *p*_*act*_ depends on the number *N* of entering ions. For a fixed probability *p*_*act*_, we plotted *r*_*max,Pact*_ (*N*) in Fig. 1D. For a sparse distribution of vesicles, characterized by a bulk located a distance *H* = 100 nm from the membrane, a vesicle is activated with a probability *p*_*act*_ = 0.8 (resp. *p*_*act*_ = 0.2), when 1200 ions are entering at a distance 100 nm (resp. 450 ions), and 340 ions at a distance 50 nm (resp. 125 ions). This result has to be compared to the 200-500 nm diameter of the AZ [41]. Consequently, a sparse distribution of vesicles at the AZ requires a high number of entering calcium ions in order to trigger fusion, which can be achieved when channels are clustered. However, when channels are co-localized with vesicles, the activation probability *pact* is significantly increased: indeed 450 ions are necessary for activation for *p*_*act*_ = 0.2 at a distance 100 nm. When the probability increases to 0.8, the distance reduces to 58 nm.

To conclude, a synapse with high release probability requires a nanometer precision of the channel location. However, this high requirement can be compensated by increasing the number of initial ions entering through VGCC clustering: with 2000 ions, the maximum distance is relaxed to 140 nm. On the contrary, in a pre-synaptic terminal crowded at its surface with vesicles (characterized by *H* = 35 nm), very few initial ions are needed for an efficient release. Indeed, 50 ions are enough to activate a vesicle with probability 0.8, wherever the channels are located in AZ (Fig. 1D).

## 5 Computing the distribution of release probability

To compute the time distribution of the release probability and to account for the calcium ions at the AZ and in the bulk, a full model of the pre-synaptic terminal is needed. The main challenge for such derivation is to account both for the stochastic regime governed by rare events of individual calcium ions arriving to a sensor binding site, and the continuous description of the calcium concentration in the bulk of the synapse [42]. The classical approach consists in using partial differential equations that often cannot take into account easily the specific AZ organization and in particular the geometry near vesicles. An important assumption of these approaches is the use of the forward rate for the calcium to sensors. This rate is often assumed and not derived (contrary to the approach described in subsection 3.1).

To compute the sensor activation, a different approach is to use Monte-Carlo or Brownian simulations to follow each ionic trajectory. But this approach is often computationally greedy to detect the rare events of calcium hitting a small target [16]. Recently, a hybrid Markov-mass action model has been developed [35], that combines a Markov chain to represent the stochastic events occurring at the AZ, with a mass-action laws model that represents calcium dynamics in the large bulk. The Markov chain and the mass-action model are coupled by the calcium ions coming from the bulk and binding to the sensor. The arrival time of such ions is Poissonian, with the rate computed taking into account the geometry of the vesicle, as discussed in subsection 3.1. This model is used to compute the time distribution of vesicular release [35] and it shows that vesicular release is triggered by the binding of calcium ions that can originate either from the bulk or from VGCC.

The distribution of release time is bimodal although it is triggered by a single fast action potential (Fig. 2). This simulation is initiated by three channels and each of them let a flow of 80 ions inside the cell during a time scale that was simulated from a Hudgkin-Huxley model [35].

**Figure 2:**
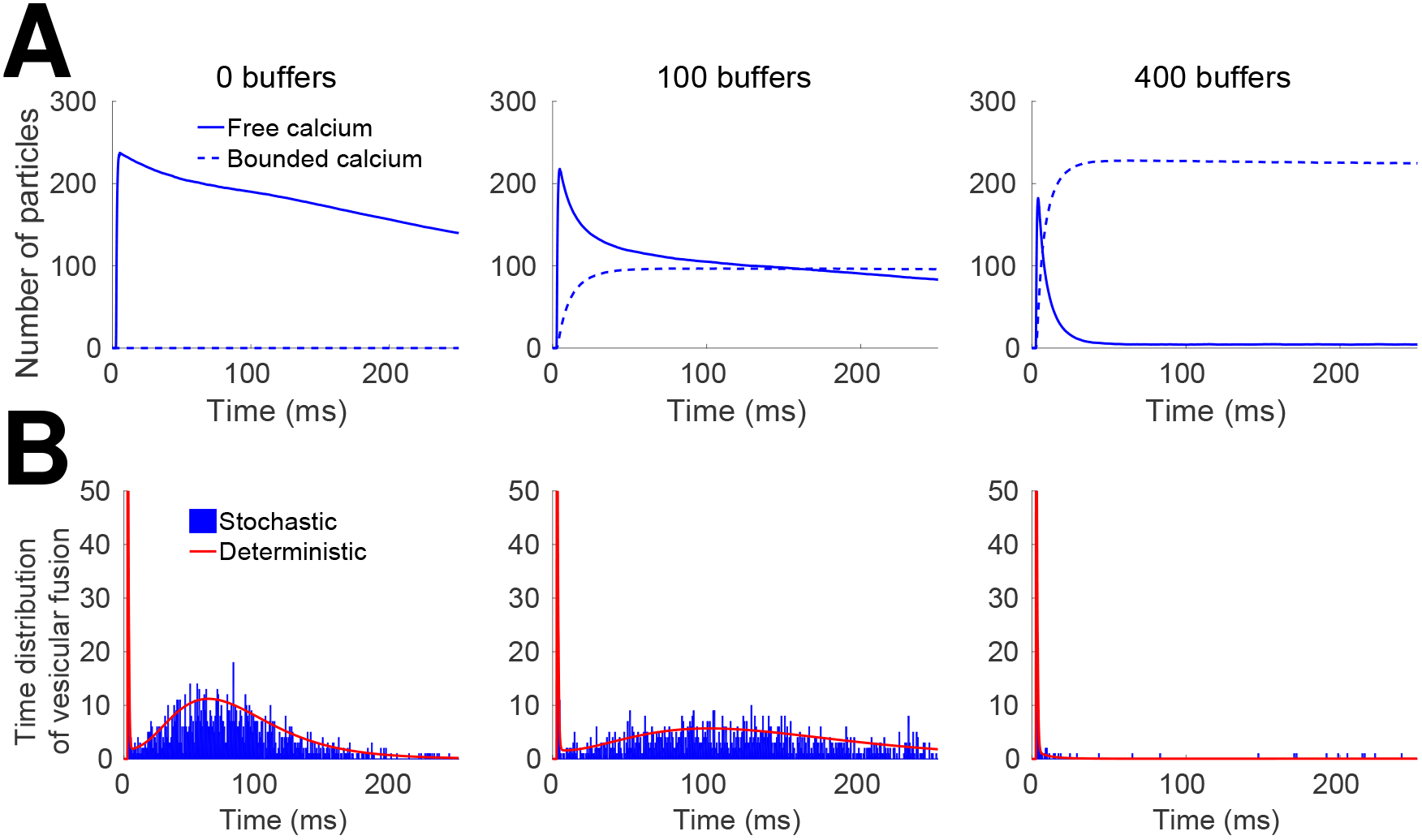
Consequences on the release probability of calcium channel location and vesicular crowding at the AZ. Calcium time course in the pre-synaptic terminal and vesicular release activation. A: Number of free (continuous) and buffered (dotted) ions for 0 (left), 100 (middle) and 400 (right) buffer sites. **B**: Histogram of vesicular release time for the stochastic (blue), and the Markov-mass action model (red) for 0 (left), 100 (middle) and 400 (right) buffer sites.

An example of specific simulation is as follows: the pre-synaptic terminal is a bulbous head of volume ≈ 1 *μm*^3^. At the AZ, we positioned eight vesicles, distributed on a square lattice of surface ≈ 0.13 *μm*^2^ (Fig. 1 and [35]), so that the distance between two neighboring vesicles is 130 nm, and each vesicle has a diameter of 40 nm. The three calcium channels are uniformly distributed over the AZ, but remained from a distance of 6 – 10 nm (we chose around 6 nm here) from every vesicle. This distance corresponds to the radius of the red ribbon (Fig. 3A) of height 1 nm the calcium ions need to reach to simulate the binding to a sensor. For each simulation, the terminal undergoes three spikes at a fixed time interval Δt = 20 – 150 ms. Once a calcium ion enters the terminal, it can either reach a vesicle with probability *p*_*s*_ (eq. 3), or enter inside the bulk with probability 1 – *p*_*s*_ according to the scheme of Fig. 3C. We already discussed in subsection 2.2 how to chose the calcium diffusion coefficient: at the AZ it is 200 *μm*^2^/*s*, and 20 *μm*^2^/*s* in the bulk to account for crowding. Inside the bulk, ions bind to buffers with a rate constant *k*_0_ = 5.6 s^−1^ and unbind with rate *k*_−1_ = 500 *s*^−1^. Calcium can be extruded by pumps with a rate *k*_*pump*_ = 0.88 *s*^−1^ or they can leave the terminal with rate *k*_*es*_ = 6.1 *s*^−1^. Finally, calcium can bind to the sensors located underneath a vesicle with a rate *k*_*S*_ = 0.3 *s*^−1^. The number of buffer molecules in the bulk varies from 0 to 1000. A calcium ion bound at the calcium sensor can unbind with a rate *k*_*u*_ = 2000 *s*^−1^ (fast unbinding) or 5 *s*^−1^ (slow unbinding). Once 5 calcium ions are bound to a calcium sensor, then the vesicle fuses with the synaptic membrane and the vesicle spot becomes free for a new vesicle coming from the recycling pool to bind with a rate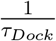(Fig.3D). An immediate refilling of vesicles at the AZ is obtained with a time *τ*_*Dock*_=0 ms, but other delay are possible such as *τ*_*Dock*_ = 50 ms, or not refilling of vesicles with *τ*_*Dock*_ = ∞ as shown in Fig. 4.

**Figure 3:**
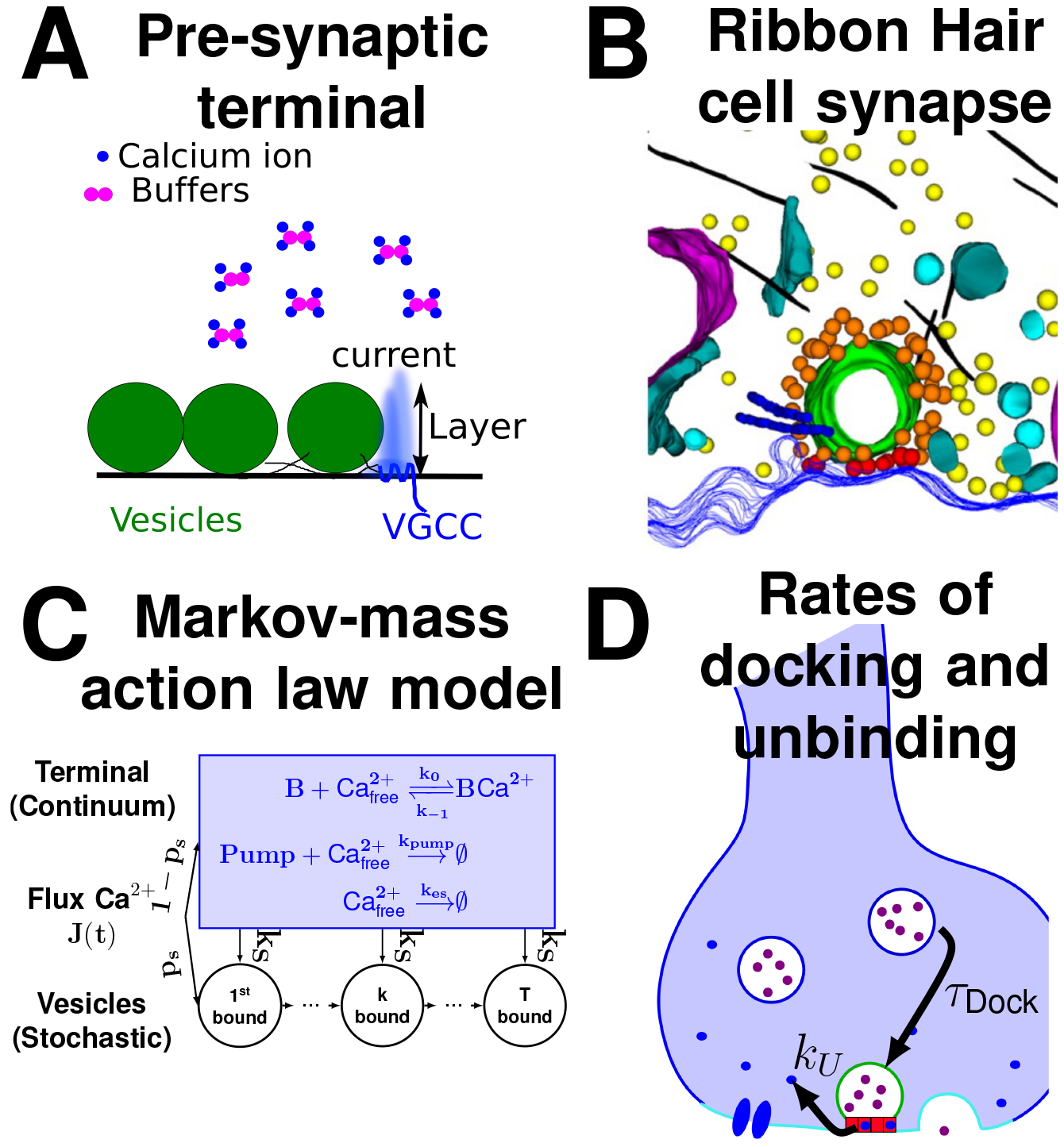
Various model of the AZ. **A:** The entering flux of calcium ions (blue) through a VGCC, could be affect by the vesicle geometry. The flux penetrates inside a layer of tens of nanometers. Buffers (purple) can bind calcium ions. **B**: Hair cell synapse containing a circular ribbon (green) that could generate a micro-domain to retain calcium between the boundary layer near the membrane and the pre-synaptic bulk (reproduced from [22]). **C**: Summary of the Markov-Mass action model, showing how the initial flux is split between the calcium vesicular sensor and the bulk [35]. **D**: Illustration of the rates: for replacing vesicles 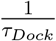 and for calcium unbinding *k*_*U*_ from the calcium sensor (red boxes).

**Figure 4:**
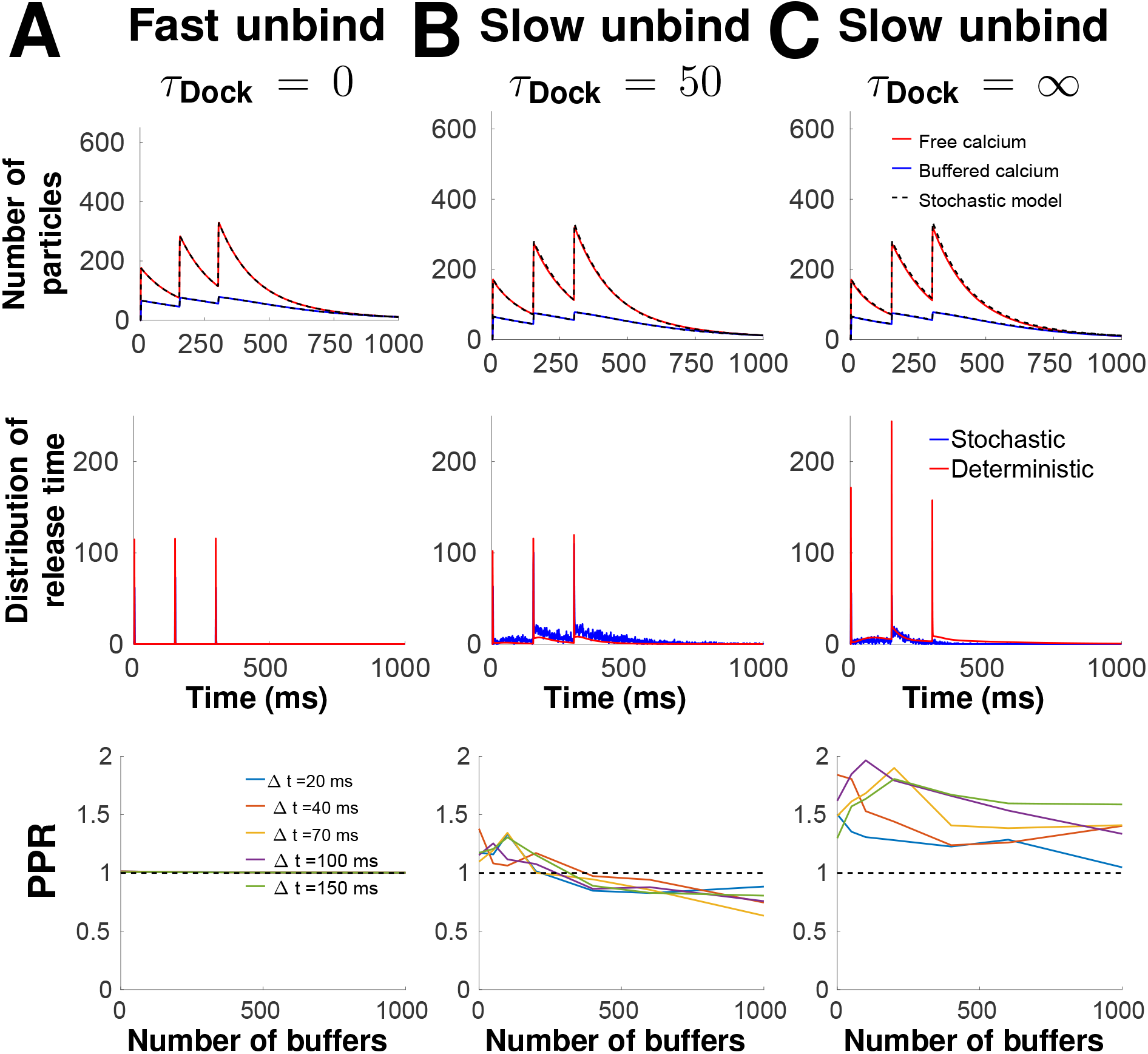
Calcium dynamic and vesicular release in the pre-synaptic terminal following three pulses. Comparison between slow unbinding from the Snare (*k*_*U*_ = 0.005 ms^−1^) with the vesicle docking time *τ*_*D*_ = 0 ms (**A**), fast unbinding (*k*_*U*_ = 2 ms^−1^) with *τ*_*D*_ = 50 ms (*B*) and fast unbinding with *τ*_*D*_ = ∞ (*C*) for 100 buffer molecules and 150 ms between two spikes. Up: Time course of free calcium (red) and buffered calcium (blue). Middle: Probability density function of the release times. Down: Paired Pulse Ratios (PPR), computed as the ratio of the number of fused vesicles during the first and the second spike.

In that simulation framework, the first peak in the time distribution of vesicular release follows a single stimulation and the second one in Fig. 2 (that is smaller in amplitude and wider) corresponds to the random arrival, over a much longer time period, of ions located in the synaptic terminal to small binding vesicular targets. To conclude, multiscale stochastic modeling approaches allow studying cellular events based on integrating discrete molecular events over long time scales from one milliseconds to seconds.

### 5.1 Asynchronous release

What defines the time lag between the arrival of an action potential and the first release of a vesicle? A possible mechanism is as follows: calcium ions flow inside a channel in less than 1ms. We saw above that calcium ions entering from VGCG located very close to a vesicle can reach key calcium sensors in less than 0.5 *μs.* Either all calcium binding sites necessary to trigger fusion are now activated and fusion does occur, or some sites are still empty. In this case, the release will then depend on the arrival to the ribbon underneath the vesicle of calcium ions that will have to travel from other places, such as the bulk or other VGCC located far away. This second arrival process has a rate constant of few seconds [35]. These random arrival times of calcium ions to the vesicular calcium sensors define the distribution of vesicular release that can be widely spread due to the two distributions of calcium sources (Fig. 2A-B). It also reveals that the distribution contains two peaks: one generated by immediate or synchronous release, which corresponds to the case where all calcium binding sites are immediately filled [42], and the second release, which is asynchronous over hundreds of milliseconds, the time scale of which is defined by the arrival of far away calcium ions.

In that case the cusp geometry underneath the vesicle defines the arrival rate (see above and [23]). In addition, it might be possible that vesicle-tethered and cytoplasmic Syntaxin1 proteins also contribute in differentially regulating synchronous versus asynchronous release kinetics [18]. In that case, the asynchronous release would be determined by the vesicle-tethered mechanism and not only by calcium arrival. The two processes could also combine together.

Under the calcium hypothesis controlling asynchronous release, increasing the concentration of calcium buffers in the bulk should reduce the amount of free calcium that can travel long distance in few milliseconds. Thus, increasing calcium buffer concentration should reduce asynchronous release, as shown in numerical simulations Fig. 2A-B [35], and experimentally [43]. Indeed, this hypothesis has received more support as buffering intracellular calcium with EGTA-AM reduced asynchronous EPSC. Asynchronous or spontaneous release involve calcium coming from the bulk or VGCC located far away from the vesicles [21, 45, 46]. The release of asynchronous vesicles was largely diminished when calcium chelation such as BAPTA [43] was used. The authors of that study concluded that asynchronous release should rely on calcium ions involving longer trajectories compared to the ones originating from VGCC located near a vesicle. Note that the number of buffer molecules such as calmodulin in the first hundred nanometers between the vesicle and the membrane is very small (of the order of a few) and thus these molecules do not affect synchronous release, as shown also experimentally in [43]. However it remains unclear whether or not the readily releasable pool organization can influence asynchronous release.

To conclude, VGCC located underneath vesicles are not the only contributor filling the calcium binding sites, required for vesicular fusion. Actually, not all VGCC are located underneath vesicles. If one or a cluster of VGCC are not close enough or are moved away from a vesicle of a distance of 10 or 20 nm, the probability to have the correct amount of calcium ions on the sensor binding sites can decrease significantly (Fig. 1D) and thus vesicular release will have to involve calcium ions coming from far away. Modeling and experiments [43] are now converging and suggest that this second source of calcium defines and regulates asynchronous release, when calcium ions generated from local VGCC is not enough. In that context, any vesicle can potentially lead to an asynchronous release as long as it does not contain enough VGCC underneath. Most likely, these vesicles are located at the periphery of the AZ, where the density of VGCC could decrease. The exact relation between VGCC distribution and vesicular organization remains unclear.

### 5.2 Simulating multiple spikes and Paired-pulse ratio

The model developed in [35, 42] can also be used to explore the short-term synaptic properties such as calcium accumulation and the distribution of time for vesicular release. First the method is consistent with any other simulation methods, second it is possible to test how the backward rate constant of the calcium ion to the sensor affects the time distribution of release, as well as the paired-pulse ratio (PPR), computed in this case as the ratio, after two consecutive spikes of the amount of fused vesicles after the second spike divided by the first one. A PPR > 1 means that the release probability is increased which is usually interpreted as short-term synaptic facilitation, while a PPR < 1 corresponds to a decrease, interpreted as short-term synaptic depression.

For fast calcium unbinding to the sensors, there is no accumulation of calcium in the sensor site and thus the release probability is independent of the spike train and of the buffer concentration (Fig. 4A, thus PPR = 1). Conversely, a slow unbinding time from the sensors is associated with a local increase in the release probability (PPR > 1) (Fig. 4B). This facilitation is due to various sources of calcium: first the ones already bound to sensors and second to calcium accumulation in the bulk following multiple spikes. This second source is diminished by increasing the amount of calcium buffers, which can lead at high buffer concentration to a decrease in the release probability PPR < 1 (Fig. 4B, bottom).

Hence, by changing the buffer concentration, a synapse can go from a facilitating state to a depressing state. To investigate the role of the readily-releasable-pool organization in the release probability, a first step is to use a vesicular replacement rate at the AZ by considering the time *τ*_Dock_ for a vesicle from the readily releasable pool of vesicles to replace a vesicle that has just fused: in the extreme case *τ*_Dock_ = 0, which corresponds to an immediate refilling of vesicles, and *τ*_Dock_ = ∞ TO which corresponds to no refilling of vesicles: Fig. 4C shows the behavior of a vesicular release with no refilling of vesicles, and a slow unbinding rate. Simulations show an increase in release probability after the second spikes, due to the calcium accumulation at the binding sites. After the third spike, the release probability is decreased, due to the lack of vesicles docked at the AZ, which would be interpreted as short-term depression. To conclude, a low unbinding rate is responsible for calcium accumulation in the sensor binding site that increases the release probability and defines short-term facilitation.

## 6 Conclusion and perspective

The lesson from modeling and numerical simulations of diffusion in the first hundred nanometers between docked vesicles and the plasma membrane is that this boundary layer is crucial for computing the vesicular release probability due to the critical position of vesicular sensors. This space is difficult to access experimentally and its role has been underestimated in short-term plasticity. However, experimental approaches using calcium chelator such as BAPTA or EGTA confirm the role of calcium ions traveling from far away compared to the ones entering directly through VGCC located underneath a vesicle to trigger release.

Another key feature relevant for short-term plasticity is the structural organization and the spatial correlation between the distribution of VGCCs and vesicles. Do vesicles contain the same amount of close VGCCs? What defines the exact location of VGCC underneath vesicles? Can this number fluctuate? What happens after vesicular fusion? How are VGCC redistributed? It is possible that VGCCs are constantly moving to find local optimal sites location [40].

Short-term facilitation is classically thought as the accumulation of calcium in the synaptic bulk (calcium hypothesis of Katz and Miledi [44]), due to various possibilities such as slow and fast buffers. But computational evidences [42] reviewed here suggest that calcium accumulation at the sensor binding sites and not in the bulk is actually the determinant effect to pre-activate vesicular release (by binding a certain fraction of the sensor sites). A similar conclusion was reached in [50] using fluctuation analysis, calcium imaging and numerical simulation analysis indicating that the residual calcium bound to the release sensors, see also [51], after the first AP could cause Paired Pulse Facilitation at Purkinje neuron synapses.

A byproduct of facilitation should be asynchronous release, because the random calcium accumulation at sensor increases the time window when a vesicular can be released, due to ions arriving at random time from the bulk, thus leading to a high variability in the calcium arrival and the vesicular release times. However, synaptic facilitation requires a low concentration of calcium buffer, suggesting that for facilitating synapses, calcium buffers should be maintained at a low level. Probably, this low concentration level can be achieved by preventing ER or mitochondria to come in too close proximity of the AZ. In contrast, synchronous release is associated with a high calcium buffer concentration, preventing calcium ions to travel from far away. More specifically, two-dimensional numerical simulations [24], modeling essentially the synaptic bulk, revealed that fixed endogenous buffers with low affinity, characterized by a low calcium-binding ratio, mixed with mobile buffers with high affinity, results in a fast AZ calcium clearance. This results in synchronous high-frequency transmission (at 200Hz). But it remains unclear how calcium fluctuations is maintained low [19, 20, 48, 49] especially near the vesicular calcium sensor.

Finally, how the rate of vesicular release can vary over 6 orders of magnitude for the same synapse [26] also remains enigmatic. The cusp geometry and rare binding events may hold the key to the solution of this spectacular modulation of the vesicular release rate [23]. Modeling vesicular trafficking and recycling at various synapses including ribbon synapses [47] should clarify the organization of the pre-synaptic terminal [22, 52] and the effect of VGCC trafficking [40].

